# Establishment of *Culex modestus* in Belgium and a glance into the virome of Belgian mosquito species

**DOI:** 10.1101/2020.11.27.401372

**Authors:** Lanjiao Wang, Ana Lucia Rosales Rosas, Lander De Coninck, Chenyan Shi, Johanna Bouckaert, Jelle Matthijnssens, Leen Delang

**Affiliations:** KU Leuven Department of Microbiology, Immunology and Transplantation, Rega Institute for Medical Research, Laboratory of Virology and Chemotherapy, Leuven, Belgium; Laboratory of Viral Metagenomics, Rega Institute for Medical Research, KU Leuven, Leuven, Belgium

## Abstract

*Culex modestus* mosquitoes are known transmission vectors of West Nile virus and Usutu virus. Their presence has been reported across several European countries, including only one larva confirmed in Belgium in 2018. Mosquitoes were collected in the city of Leuven and surroundings in the summer of 2019 and 2020. Species identification was performed based on morphological features and partial sequences of the mitochondrial cytochrome oxidase subunit 1 (COI) gene. The 107 mosquitoes collected in 2019 belonged to eight mosquito species: *Cx. pipiens* (24.3%), *Cx. modestus* (48.6%), *Cx. torrentium* (0.9%), *Culiseta annulata* (0.9%), *Culiseta morsitans* (0.9%), *Ae. sticticus* (14.0%), *Ae. cinereus* (9.3%) and *Anopheles plumbeus* (0.9%), suggesting the presence of an established *Cx. modestus* population in Belgium. Collection of *Cx. modestus* mosquitoes at the same locations in 2020, confirmed the establishment in the region.

Haplotype network analysis of the COI sequences for *Cx. modestus* showed that the Belgian population is rather diverse, suggesting that it may have been established in Belgium for some time. The Belgian *Cx. modestus* population was most closely related to populations from the UK and Germany. Characterization of the virome of the collected mosquitoes resulted in the identification of at least 33 eukaryotic viral species. Nine (near-) complete genomes belonging to 6 viral species were identified, all of which were closely related to known viruses.

In conclusion, we here report the presence of *Cx. modestus* in the surroundings of Leuven, Belgium. As this species is known to be a vector of several arboviruses, the implementation of vector surveillance programs monitoring this species is recommended.

**Importance:** *Culex modestus* is a mosquito species that plays a role as a ‘bridge’ vector, being able to transmit pathogens between birds, as well as from birds to mammals, including humans. In Belgium, this mosquito species was considered absent, until the finding of one larva in 2018 and subsequent evidence of a large population in 2019-2020 described here. We collected mosquitoes in the summer of 2019 and 2020 in the city of Leuven and surroundings. The mosquito species was identified by morphological and molecular methods, demonstrating the presence of *Cx. modestus* in this region. The ability of mosquitoes to transmit pathogens can depend on several factors, one of them being their natural virus composition. Therefore, we identified the mosquito-specific viruses harboured by Belgian mosquitoes. As *Cx. modestus* is able to transmit viruses such as West Nile virus and Usutu virus, the establishment of this mosquito species may increase the risk of virus transmission in the region. It is thus advisable to implement mosquito surveillance programs monitoring this species.

## Introduction

The *Culex modestus* mosquito species was described for the first time by Eugenio Ficalbi (1889) in northern Italy (1) and is considered a rare species. In Europe, this species is distributed mainly in southern and central European countries. Field collections have reported the presence of *Cx. modestus* in France, Spain, Portugal, Germany, Romania, Serbia and Czech Republic; and more recently, also in more northern countries such as the United Kingdom (UK), Denmark and Sweden (2–5). In Belgium, this mosquito species was thought to be likely present given its occurrence in nearby countries (6). Up to now, only one larva has been found in 2018 and identified as such through molecular methods (7). Recent field studies in the UK have confirmed two characteristics of *Cx. modestus*: (i) its ornithophilic habit, i.e. feeding on resident and migratory bird species (8), and (ii) its mammalophilic and anthropophilic feeding behaviour, showing that *Cx. modestus* is also a major human-biting mosquito species similar to *Cx. pipiens* (9). Thus, *Cx. modestus* could play a role in nature as a ‘bridge’ vector, being able to transmit pathogens between birds in an enzootic cycle, as well as from birds to mammals, including humans, in an epizootic/epidemic cycle.

Previous studies on different *Cx. modestus* populations in Europe revealed that this species can act as a carrier of different pathogens, and likely is able to transmit these pathogens as well. In the south of France, *Cx. modestus* mosquitoes have been found to serve as amplifying vectors for seasonal West Nile virus (WNV), introduced by migratory birds (10). *Cx. modestus* mosquitoes collected in the Danube Delta region (border of Romania and Ukraine) were positive for *Plasmodium sp*. lineage Donana03 (avian malaria) (11). In addition, a prevalence of 5.1% of *Trypanosomatids* was detected in the gut of *Cx. modestus* collected in the Czech Republic between 1998 and 2002 (12). Furthermore, *Cx. modestus* is the vector and reservoir of Lednice virus (LEDV), a rare bunyavirus that causes viremia in wild birds. During the last sixty years, various European countries have reported the presence of LEDV in their *Cx. modestus* mosquito populations (13). Besides LEDV, also the Tahyna virus (TAHV) has been isolated from *Cx. modestus* in Czechoslovakia and in France (14).

Mosquito surveillance in the UK started focusing on *Cx. modestus* due to its confirmed establishment and important role in the transmission of WNV and Usutu virus (USUV) (15). The role in WNV transmission in Europe was demonstrated by the detection of WNV in this mosquito species during an outbreak in the Sardinia region (Italy) in 2011 (16). During this outbreak, the circulating virus strains belonged to lineage 1. It was the first report of an Italian WNV strain that caused clinical signs in the affected birds. The mosquito survey carried out in this area revealed that these virus strains were found in *Cx. modestus* mosquitoes. During the mosquito seasons 2015 and 2016, WNV lineage 2 has also been detected in *Cx. modestus* mosquitoes collected in the Lednice-Valtice Area (in southern Moravia) (17, 18). Regarding the vector competence of *Cx. modestus* for WNV, this mosquito species was found to be competent to transmit WNV experimentally. More than 90% of *Cx. modestus* mosquitoes developed a disseminated infection 14 days after an infectious WNV bloodmeal (19). Moreover, it is considered an extremely efficient vector, given that the disseminated infection and the transmission rates reached 89.2% and 54.5% respectively, after 14 incubation days (20). The USUV has also been detected in field collected *Cx. modestus*, likely co-circulating with WNV (21). The USUV is another arbovirus with African origin that is principally transmitted by *Culex* mosquitoes. This virus belongs to the genus *Flavivirus*, as well as dengue, yellow fever, Zika, Japanese encephalitis and WNV (22). The virus is maintained in an enzootic cycle between ornithophilic mosquitoes and birds. In Europe, USUV was found by retrospective analysis of archived tissue samples from bird deaths in the Tuscany region of Italy in 1996 (23). In 2001, USUV-associated death of blackbirds was reported in Austria (24), Germany and the Netherlands (25, 26). In 2016, numerous wild birds, mainly Eurasian blackbirds (*Turdus merula*), were affected by a USUV outbreak in Belgium in the provinces of Limburg, Antwerp and Flemish Brabant (27). In 2017, the virus further spread to the west and by the summer of 2018, the whole country was affected (28). Despite the recent USUV outbreaks, it is not known which mosquito species are the vectors of USUV in Belgium. To gain insight about which mosquito species might carry clinically relevant viruses, we collected field mosquitoes using BG-sentinel traps in the city of Leuven and its surroundings in three different environment types (urban, peri-urban and wetland areas).

To unravel the high diversity of mosquito-specific viruses (MSVs) harbored by Belgian mosquitoes, we performed a metagenomic sequencing approach using the Novel enrichment technique of VIRomes (NetoVIR) protocol (29). The study of viral diversity in mosquitoes is important since MSVs have the potential to modulate the vector competence of mosquitoes for different arboviruses (30). The virome of tropical mosquito species such as *Aedes aegypti* has been studied extensively. On the other hand, knowledge on the viral diversity in mosquitoes from more temperate regions is still scarce but increasing. For instance, a recent virome study identified novel RNA viruses in Swedish mosquitoes (31). However, the virome of mosquitoes from Western Europe, including Belgium, has not yet been studied. Therefore, we provide a first glance into the virome of mosquitoes collected in Belgium.

## Material and Methods

### Ethics statement

Permits for peri-urban and wetland mosquito field collections were obtained from the security responsible of KU Leuven. Permits for field collections in urban habitats were obtained from the landowners.

### Mosquito collections

Adult mosquitoes were trapped with the BG-Sentinel traps (BioGents GmbH, Germany), which were baited with BG-lure (BioGents GmbH, Germany) and containing around 2 kg of dry ice in the isolated box for CO_2_ production. Two traps were rotated in three different habitat types (urban (N 50°52’41, E 4°41’21), peri-urban (N 50°51’, E 4°41’), and water reservoir wetlands (N 50°51’, E 4°40’), in Leuven and surroundings (Supplementary Figure S1).

The parameters to determine each trap location in these habitats were similar to what is described by Mayi et al. (2020) (32). We followed these criteria and the advice of Prof. dr. ir. Raf Aerts and his team at the Division Ecology, Evolution and Biodiversity Conservation, University of Leuven (KU Leuven) on the selection of mosquito collection sites representing different habitat types.

Collections were performed from August to the beginning of October in 2019, when the weather was good, avoiding strong wind or heavy rain. Every 24 hours, the traps were emptied and repositioned between sunrise and sunset of the next day. Mosquitoes were individually stored at −80°C until species identification. A second collection was performed in August of 2020 in the same geographic locations as described above to confirm the presence of certain species.

### Species identification, sample preparation and DNA sequencing

All collected mosquitoes were identified using morphological characters (33). Individual thoraces were removed using forceps for molecular identification and homogenized in 100 μL of phosphate-buffered saline (PBS) using tubes with 2.8 mm ceramic beads with a Precellys Evolution homogenizer. Sample preparation was performed by lysing the homogenate at 100°C for 10 minutes (34). Tissue debris was removed by centrifugation at 12 000 rpm for 3 minutes, and 50 μL of supernatant was collected into a new tube. A 710 bp region of the cytochrome oxidase subunit I (COI) mitochondrial gene was the target for amplification by polymerase chain reaction (PCR) using previously reported primers (35). Presence of the PCR product was checked on a 2% agarose gel by gel electrophoresis. DNA was purified with the Wizard^®^ SV Gel and PCR Clean system (Promega). The DNA concentration of amplicon was measured by NanoDrop (ThermoFisher), after which samples were sent for Sanger sequencing to Macrogen Europe.

### Mosquito sequence analysis and phylogeny

Sequences were edited and assembled with Bioedit version 7.2.5 (36) to obtain a single consensus sequence per individual mosquito. Through the BLAST tool, the generated COI sequences were compared to the NCBI database. Reference COI sequences for all mosquito species considered were selected according to Versteirt and colleagues (37), which employed reference sequences that were registered in the Barcode of Life Data (BOLD) Systems, and downloaded from GenBank. For phylogenetic analysis, the COI sequences generated in the study and the reference sequences were aligned with MAFFT v7.471 (38) using the G-INS-I option. The resulting alignment was trimmed by trimAl v1.4.rev15 (39) on gappyout setting and phylogenetic informative regions of the alignment were selected with BMGE v1.12 (40) for phylogenetic inference. Maximum-likelihood (ML) trees were constructed using IQ-TREE v2.0.3 (41) with automatic selection of the best nucleotide substitution model and 1000 ultrafast bootstrap replicates. Finally, trees were visualized using FigTree v1.4.4.

### Haplotype network

Haplotype inference and nucleotide diversity were calculated in ARLEQUIN, version 3.5.2.2 (42). The population genetic data was analyzed using the median-joining (MJ) network algorithm in PopART, version 1.7 (43, 44). The COI sequences for *Cx. modestus* included in the haplotype network were retrieved from the NCBI database. These sequences were selected based on the specimen’s country of origin and the length of the COI fragment (35).

### Pool design, sample preparation and sequencing for virome analysis

The samples of mosquito abdomens were grouped in pools for sequencing according to the morphological identification of mosquito species by key points and sample location (urban, peri-urban and wetlands). Abdomens were homogenized in 600 μL of PBS with 2.8 mm ceramic beads with the MINILYS tissue homogenizer, including a negative control (blank tube with PBS).

All pool samples followed the Novel enrichment technique of VIRomes (NetoVIR) sample preparation protocol optimized for viral metagenomics (45, 46). In brief, after homogenization, samples went through a centrifugation and filtration step to remove pro- and eukaryotic organisms and large organic debris. Next, a nuclease treatment (employing benzonase and micrococcal nuclease) was applied to remove free floating nucleic acids. Nucleic acids were extracted with the QIAamp Viral RNA mini kit (QIAGEN) to be further randomly amplified using a modified Whole Transcriptome Amplification 2 (WTA2) kit procedure (Sigma-Aldrich). The products were purified, and libraries were prepared using the NexteraXT Library Preparation kit (Illumina). Sequencing of the samples was carried out on a NextSeq 500 High Throughput platform (Illumina) for 300 cycles.

### Bioinformatic analysis and identification of eukaryotic viruses

Quality and adapter trimming on raw paired-end reads was performed using Trimmomatic v0.39 (47). Next, contamination of samples was removed with Bowtie2 v2.3.4 (48) by mapping trimmed reads to a set of contigs present in the negative controls (reagent contamination). Remaining reads were *de novo* assembled into contigs using metaSPAdes v3.13.0 (49). To remove redundancy in the data, contigs were filtered on a length of 1000bp and subsequently clustered at 95% nucleotide identity over 80% of the length using Cluster-Genomes (https://bitbucket.org/MAVERICLab/docker-clustergenomes). All contigs were classified by DIAMOND (50) against NCBI’s nr database (downloaded on 27 October 2020) on sensitive mode for taxonomic annotation. KronaTools (51) was used to parse the DIAMOND output file and find the least common ancestor for each contig (based on the best 25 DIAMOND hits). Contigs annotated as eukaryotic virus were retrieved using an in-house Python script. Pool magnitudes were obtained by mapping the trimmed and decontaminated reads to these eukaryotic viral contigs with BWA-MEM2 (52, 53). The resulting abundance table was further used for ecological analysis in R using the phyloseq (54), metagenomeSeq (55), vegan (56) and ComplexHeatmap (57) packages.

### Recovery and phylogenetic analysis of (near-) complete meta-assembled genomes

To recover full eukaryotic viral genomes in the mosquito pools, viral species were selected based on the level of genome completion after metagenomic *de novo* assembly. If a viral genome was not yet fully complete after assembly, the reads from the mosquito pool were mapped to a selected reference sequence (based on the annotated species by DIAMOND and Krona tools) with BWA-MEM2 (52, 53). The consensus sequence was subsequently retrieved with samtools and bcftools (58). For phylogenetic analysis, relevant reference complete genome sequences were chosen after BLASTn of the metagenomic assembled genomes (MAGs) and subsequently downloaded from GenBank. Alignment, trim, model selection, construction and visualization of phylogenetic trees was done as previously described for mosquito COI sequences.

## Results

### Mosquito species detected in Leuven, Belgium

A total of 107 mosquito specimens was collected in three distinct locations in Leuven in the summer of 2019. According to the DNA barcodes generated and morphological features, these mosquitoes belonged to eight mosquito species: *Cx. pipiens* (24.3%), *Cx. modestus* (48.6%), *Cx. torrentium* (0.9%), *Culiseta annulata* (0.9%), *Culiseta morsitans* (0.9%), *Ae. sticticus* (14.0%), *Ae. cinereus* (9.3%) and *Anopheles plumbeus* (0.9%) (Figure 1A). Surprisingly, *Cx. modestus* accounted for ~50% of all collected mosquitoes in three different breeding sites. *Cx*. species were predominant in urban and peri-urban areas, whereas specimens found in the water reservoir wetlands belonged mostly to the genus *Aedes* (Figure 1B).

**Figure 1.**
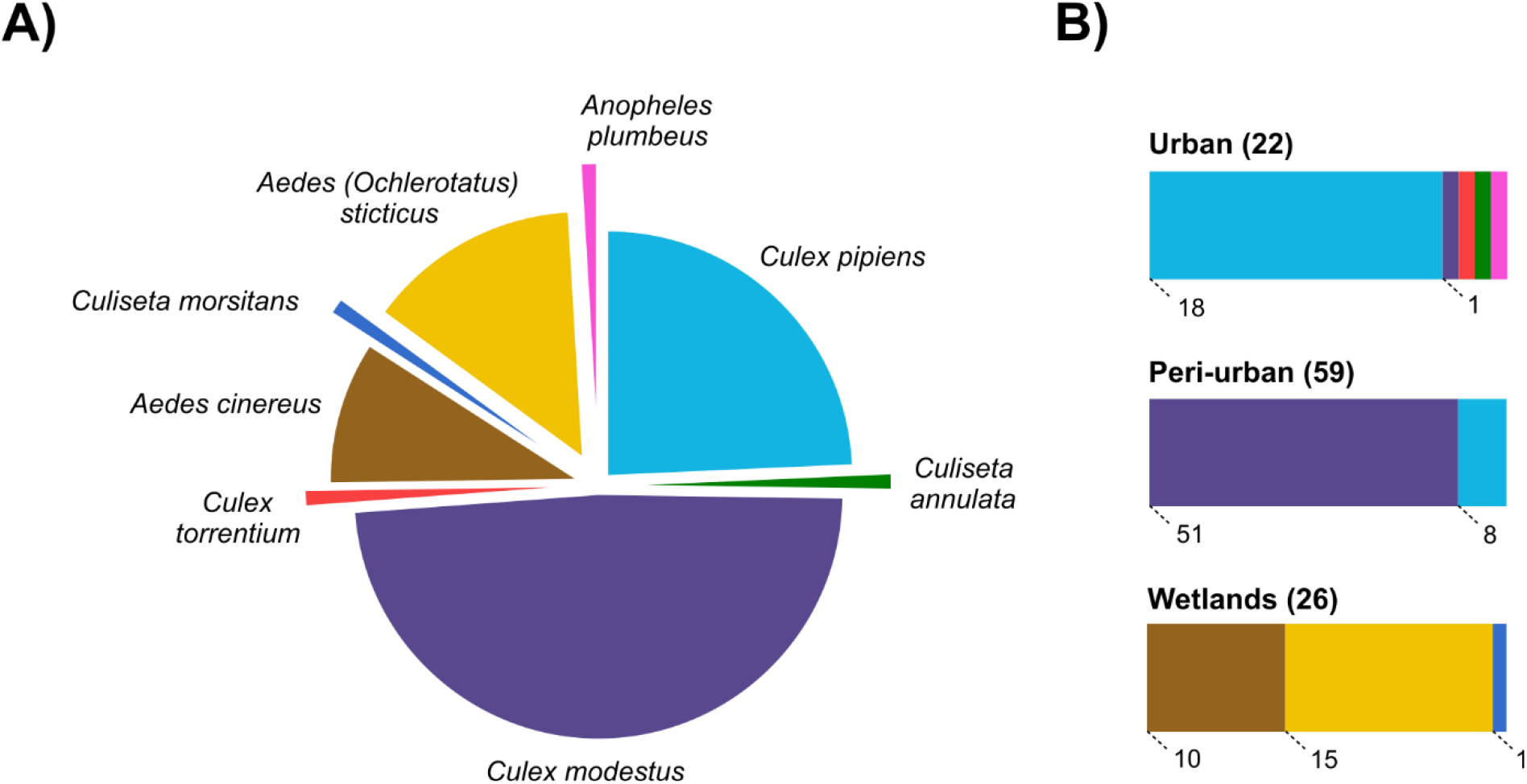
Mosquito species collected in Leuven, Belgium in 2019. A) Distribution of mosquito species captured during the summer of 2019 across all locations sampled in Leuven. B) Distribution of mosquito species across habitat types in Leuven. Mosquito species are marked in different colors. The number of specimens is indicated in the bar chart.

### Establishment of *Cx. modestus* in Leuven, Belgium

A ML tree was built from the *Cx. modestus* COI barcodes obtained in Leuven and COI sequences of 20 other Culicid species described in (6). *Cx. modestus* barcodes from Leuven clustered with two reference *Cx. modestus* sequences that were included (KJ401305, MK971991). All sequences for *Cx. modestus* fell within one large well-supported monophyletic cluster, separated from other mosquito species, which suggests that they belong to the same species (Figure 2). To find out whether *Cx. modestus* is established in the region, field collections were performed in the summer of the consecutive year (2020) using the same geographic locations as previously. Again, *Cx. modestus* mosquitoes were retrieved (Figure 2), confirming the establishment of this mosquito species in the area of Leuven.

**Figure 2.**
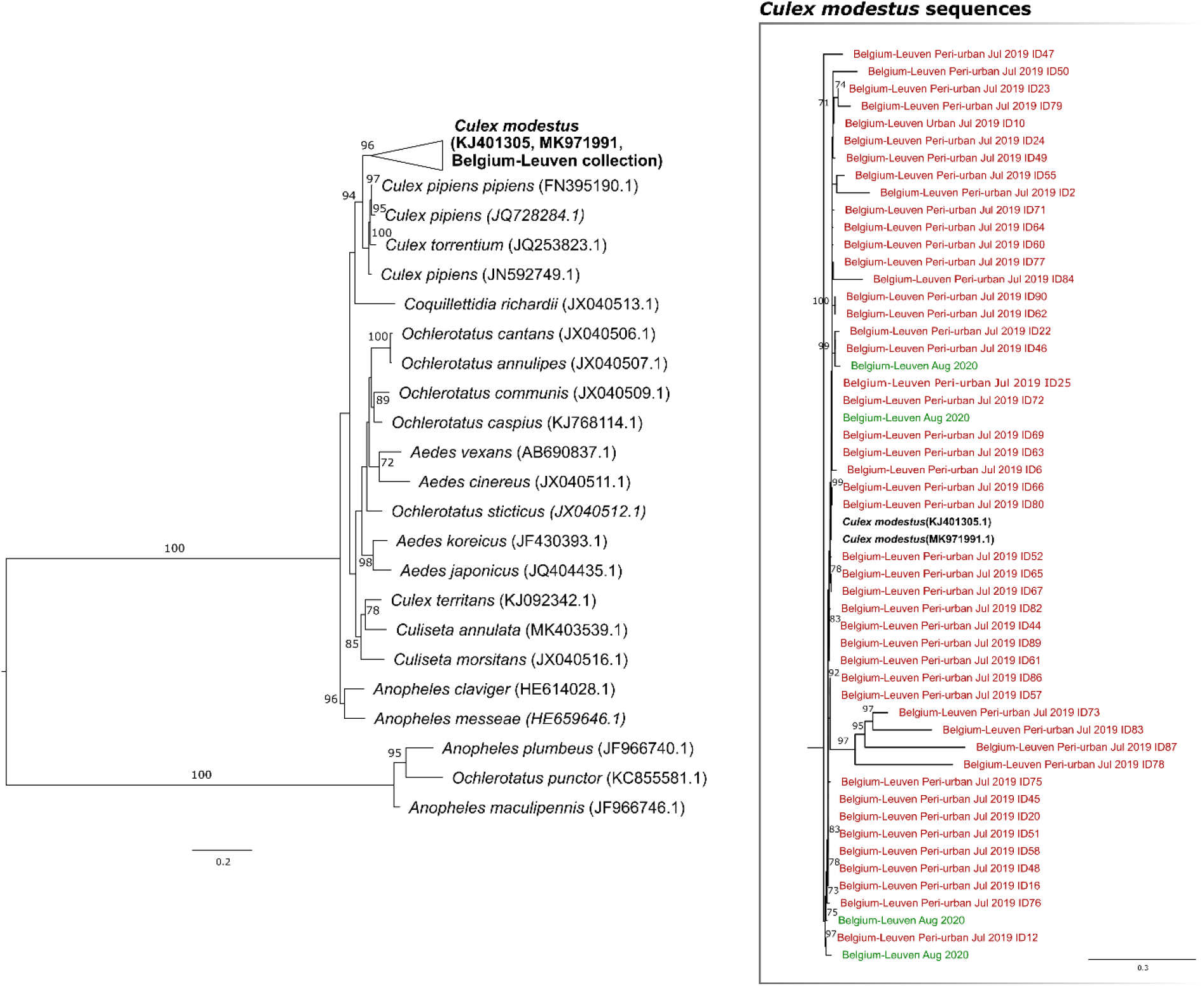
ML tree of the COI sequences of 21 Culicid species. Sequences derived from mosquitoes collected in Leuven are collapsed with the reference sequences for *Cx. modestus*. The collapsed branched is expanded in the panel on the right. Bootstrap values above 70% are shown above the branch points.

### Haplotype network of *Cx. modestus* mosquitoes

The dataset analyzed for haplotype inference was constructed employing 184 *Cx. modestus* partial COI sequences retrieved from NCBI corresponding to eight European countries, and including 40 partial high-quality COI sequences obtained from the molecular identification of field collected mosquitoes in Leuven (Supplementary Table S2, S3, S4). 4 partial COI sequences from mosquitoes collected during the summer of 2020 were included as well.

Among the 228 COI sequences (639 bp), 97 haplotypes were found. The majority of haplotypes (88) were present only in the country of origin, while only 9 haplotypes were shared by two or more countries. Haplotype diversity ranged from 0.8182 in Spain to 1.000 in Denmark, Portugal, Serbia and Sweden (Table 1). This analysis revealed that haplotype diversity in Belgium was the second highest (0.9852) of all countries screened, followed by the U.K. (0.9252) and Germany (0.9013). Nucleotide diversity estimations ranged from 0.0058 in Spain to 0.0270 in Belgium. Belgium exhibited a nucleotide diversity of 0.0270, which can be considered moderate, but which is the highest in all included European countries.

**Table 1.** Haplotype and nucleotide diversity of *Cx. modestus* from 9 countries in Europe.

| Population | *n* | Number of haplotypes | Haplotype diversity | Nucleotide diversity |
| --- | --- | --- | --- | --- |
| Belgium | 44 | 33 | $0.9852 \pm 0.0082$ | $0.0270 \pm 0.0136$ |
| Denmark | 7 | 7 | $1.0000 \pm 0.0764$ | $0.0174 \pm 0.0103$ |
| France | 28 | 11 | $0.8598 \pm 0.0462$ | $0.0069 \pm 0.0039$ |
| Germany | 42 | 17 | $0.9013 \pm 0.0278$ | $0.0113 \pm 0.0060$ |
| Portugal | 2 | 2 | $1.0000 \pm 0.5000$ | $0.0065 \pm 0.0073$ |
| Serbia | 4 | 4 | $1.0000 \pm 0.1768$ | $0.0182 \pm 0.0125$ |
| Spain | 22 | 8 | $0.8182 \pm 0.0586$ | $0.0058 \pm 0.0034$ |
| Sweden | 5 | 5 | $1.0000 \pm 0.1265$ | $0.0085 \pm 0.0058$ |
| UK | 74 | 28 | $0.9252 \pm 0.0176$ | $0.0107 \pm 0.0057$ |

### Mitochondrial DNA genealogy of *Cx. modestus*

The median-joining network displayed the ancestry of *Cx. modestus* mosquitoes (Figure 3), where two lineages were visualized separated by 1 mutation step. Haplotypes from Spain and Portugal were found uniquely in lineage I, while haplotypes from Germany, the UK, Belgium and Sweden predominated in lineage II. Haplotypes from France, Serbia and Denmark were scattered across both lineages. The majority of haplotypes that were found in Belgium were located in between of three central haplotypes of lineage II, which contain samples from several countries: one is shared by Belgium, the UK and Serbia (Figure 2, “3”), another one is shared by Belgium, the UK and Sweden (Figure 2, “2”), and the biggest one is shared by Belgium, Germany, the UK, France and Sweden (Figure 2, “1”). Haplotypes found in mosquitoes collected in Leuven during the summer of 2020 were observed in both lineage I (1 haplotype) and lineage II (3 haplotypes).

**Figure 3.**
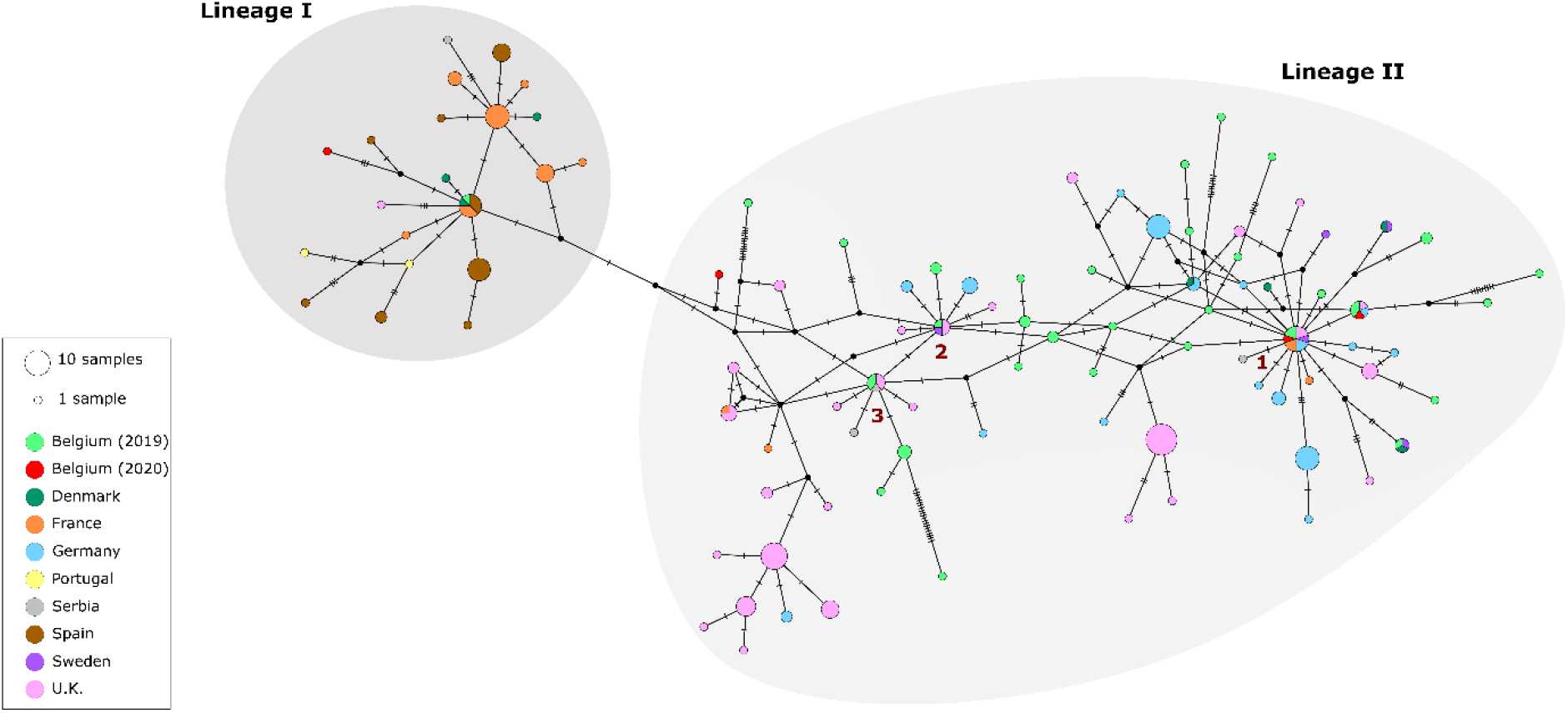
Median-joining network constructed with 228 COI sequences of *Cx. modestus* from 9 countries in Europe. Each circle represents a haplotype. The size of the circle corresponds to the number of specimens sharing that specific haplotype. Each country is represented by a color described in the legend. Mosquito collections in Belgium are separated per year to visualize the allocation of haplotypes in the network.

### A peek into the virome of Belgian mosquitoes

We characterized the virome of 107 mosquitoes’ abdomens, divided into eight pools according to their morphological identification and representing the three different habitat types mentioned before. A total of 44,002,358 reads were obtained from all mosquito pools. Most reads (21,602,296; 49.1%) belonged to the urban group. Mosquitoes collected in peri-urban and wetland areas generated 13,891,285 (31.6%) and 8,508,777 (19.3%) reads, respectively.

In all pools, the proportion of reads mapping to the order Diptera ranged from 40.8 to 77.7%. Regarding the bacterial reads, the wetland samples had a higher mean proportion (3.83%), followed by the urban samples with 2.03%, while the peri-urban samples presented less than 1% of reads mapping to bacteria. The viral component was more variable, with an observable ascending trend when moving from the wetlands to peri-urban and urban areas. Wetland samples gathered a low proportion of viral reads (<2%), whereas viral reads in peri-urban areas accounted for 1.28 – 7.19%. Lastly, reads mapping to the viral component composed 7.45 – 44.69% in the urban samples.

After filtering the viral reads for eukaryotic viral species, the relative abundances in the mosquito pools are shown in Figure 4 per viral family. While the *Cx*. pools in the urban area were completely dominated by one viral family (*Mesoniviridae* and *Iflaviridae* for pool 1 and pool 2 respectively), the mosquito pools from the peri-urban and wetland habitats seemed to have a higher viral diversity. The peri-urban samples contained mostly viral reads from a Negev-related virus, namely Yongsan negev-like virus 1, and from the *Totiviridae* family, with Culex inatomii totivirus being the most abundant viral species. In the wetlands *Ae. cinereus* pool on the other hand, an unclassified Bunya-like virus was most abundant.

**Figure 4.**
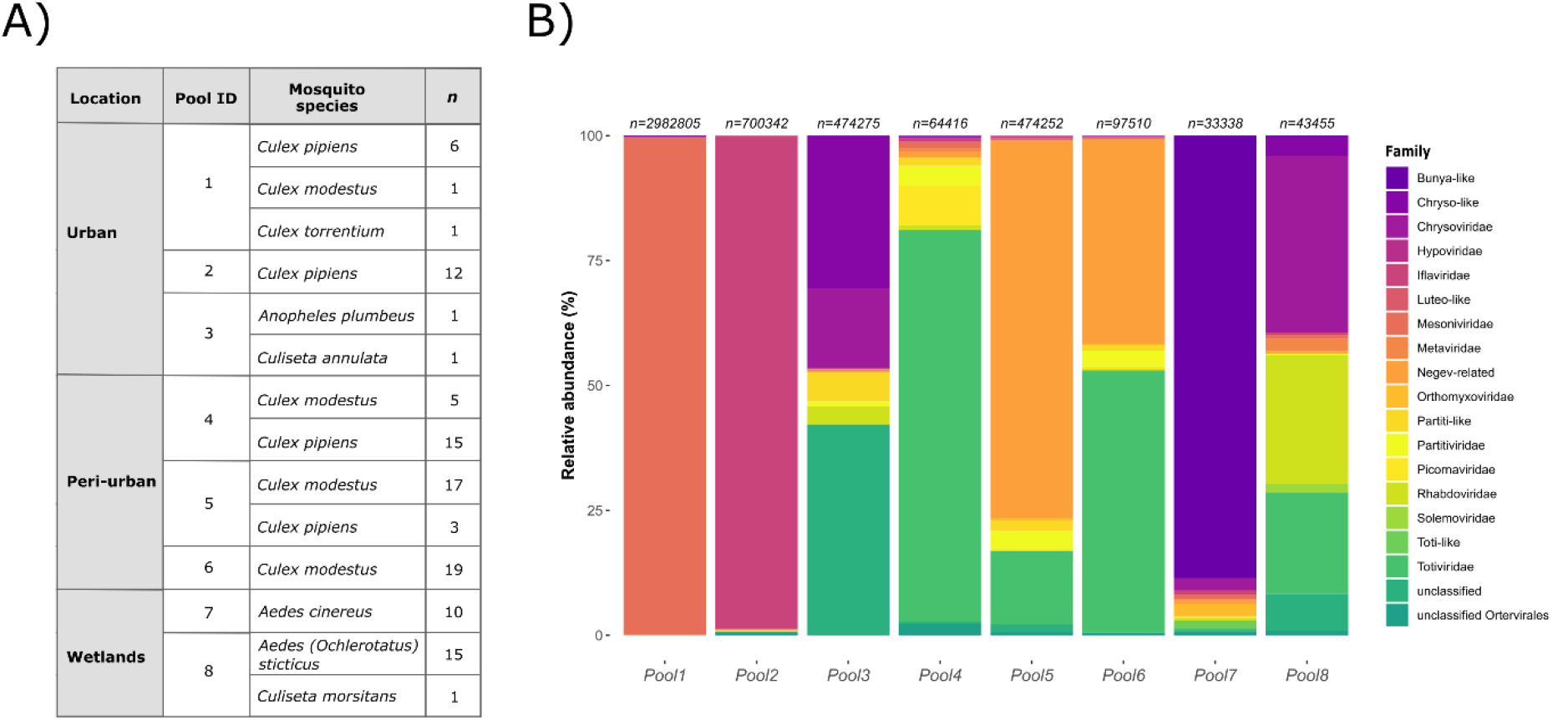
Summary information and viral composition of sequenced samples. A) Location, mosquito species and number of specimens present in each of the sequenced pools. B) Barplots representing the abundance of reads belonging to distinct viral families per pool. The number of eukaryotic viral reads per pool is given on top of each bar.

### Comparing the eukaryotic virome across habitat type and mosquito genus

To compare the eukaryotic virome of our samples, we mapped all trimmed and decontaminated reads back to the selected viral contigs, extracted the abundance table and subsequently constructed a heatmap with the normalized counts for each viral species on a log_2_ scale (Figure 5). In total, 33 eukaryotic viral species could be detected across all samples (a viral species was considered present if it had at least one contig >1000 bp and if more than 500 reads map to it). According to the Bray-Curtis distance matrix, the eukaryotic viromes of the *Cx*. mosquito pools clearly clustered together per habitat type. However, except for the peri-urban *Cx*. pools, each remaining pool had a more unique viral composition and only a small number of viruses were significantly shared between samples. Nevertheless, the peri-urban mosquito pools had a majority of viruses in common, such as Culex inatomii totivirus and Yongsan negev-like virus 1 which were shared with high abundance, while Ista virus, Sonnbo virus and Fitzroy Crossing toti-like virus 2 were common in lower abundance.

**Figure 5.**
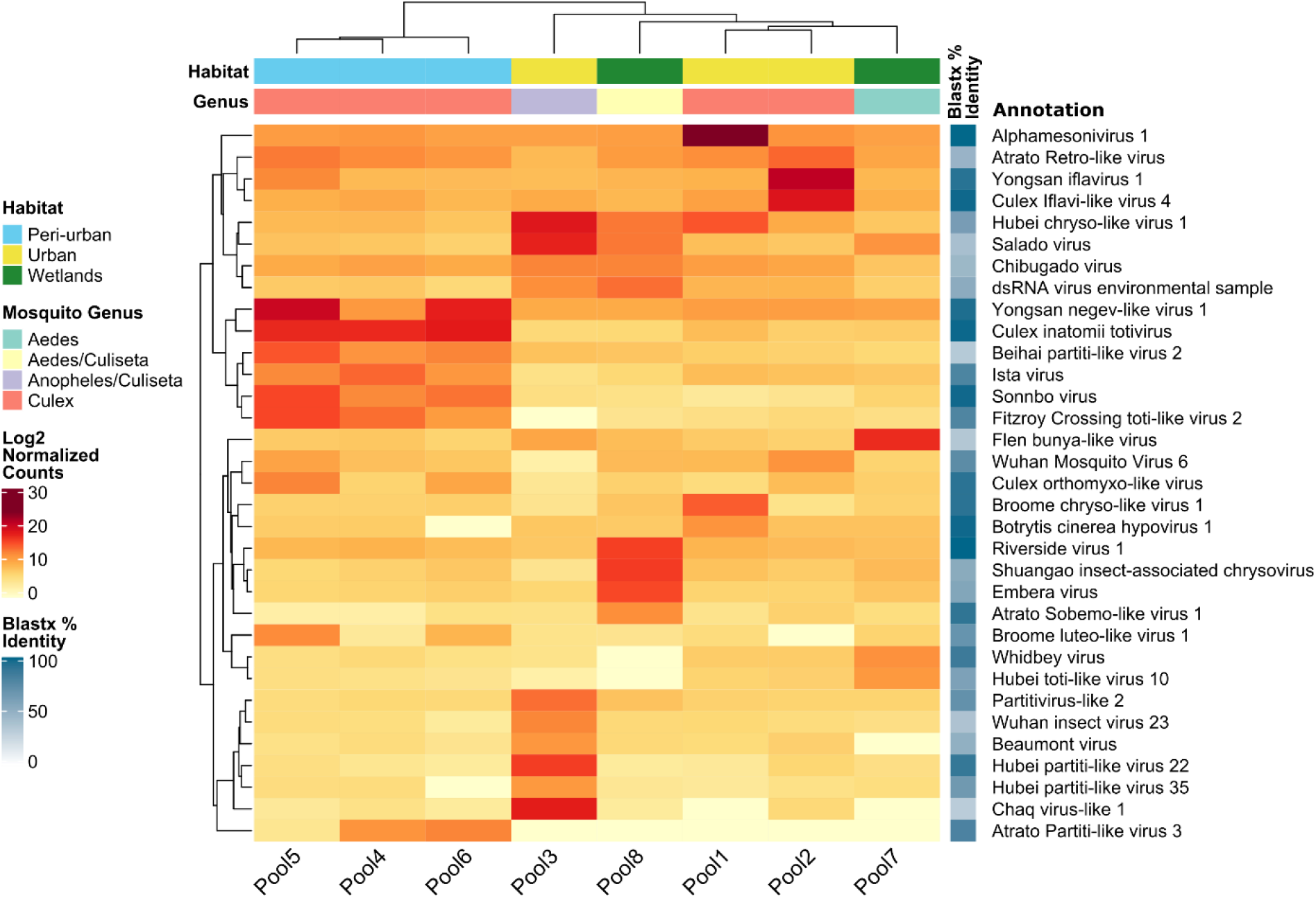
Heatmap of normalized read counts for eukaryotic viruses. The heatmap shows the normalized count on log_2_ scale of reads mapping to the assembled contigs of each eukaryotic virus. Next to the taxonomic annotation, obtained by DIAMOND and KronaTools, the average BLASTx identity for all contigs representing a viral species is depicted by the shaded blue boxes. Hierarchical clustering of the columns is based on the Bray-Curtis distance matrix calculated from the normalized read counts.

### Recovery of (near-) complete meta-assembled genomes

In total we managed to recover 9 (near-) complete genomes of 6 viral species in our metagenomic data. These viral species belong to the following families: *Totiviridae* (Culex inatomii totivirus in pool 4, 5 and 6), *Iflaviridae* (Yongsan iflavirus 1 and Culex iflavi-like virus 4 in pool 2), *Mesoniviridae* (Alphamesonivirus 1 in pool 1), *Rhabdoviridae* (Riverside virus 1 in pool 8) and unclassified Negev-related viruses (Yongsan negev-like virus 1 in pool 5 and 6), and their phylogenetic relatedness to closely related reference strains is shown in Figure 6 (Supplementary Table S5).

**Figure 6.**
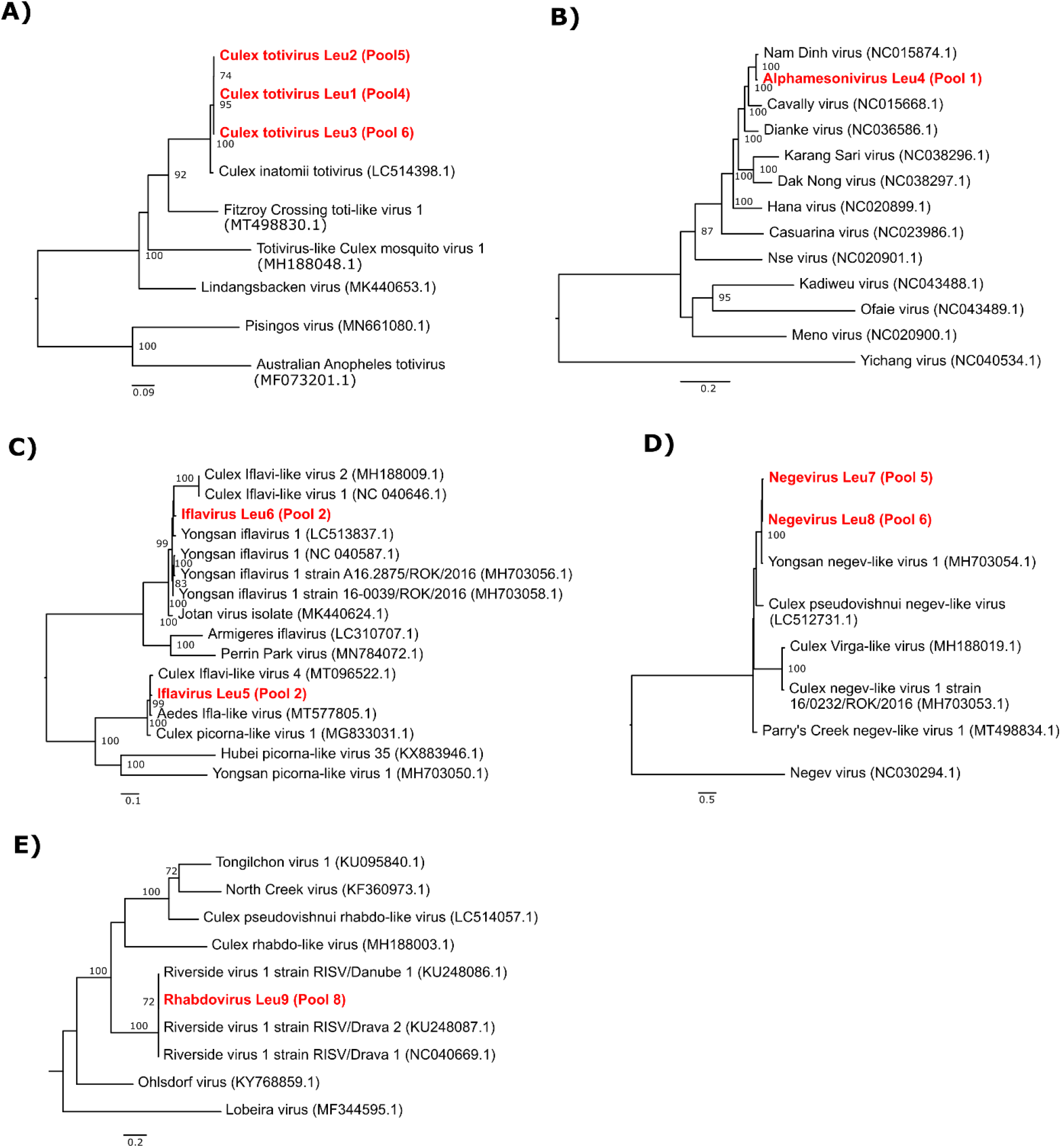
(Near-) complete meta-assembled genomes identified in mosquitoes collected during the summer of 2019. Bootstrap support values are shown next to the nodes. Complete MAGs are coloured in red. A) Midpoint-rooted ML tree of all complete genomes related to Culex inatomii totivirus, selected after BLASTn. B) Midpoint-rooted ML tree of all *Mesoniviridae* family members. C) Midpoint-rooted ML tree of all complete genomes related to our Yongsan iflavirus 1 and Culex Iflavi-like virus 4 genomes, selected after BLASTn. D) ML tree of all complete genomes related to Yongsan negev-like virus 1, selected after BLASTn. Negevirus was used as the outgroup. E) Midpoint-rooted ML tree of all complete genomes related to the recovered Riversidevirus 1.

### dsRNA viruses

#### Totiviridae

This family of dsRNA viruses are known to infect fungi, plants and invertebrates. In this study, we found Culex totivirus Leu1, Leu2 and Leu3 (98.3% average BLASTx identity with Culex inatomii totivirus; LC514398.1) in all peri-urban mosquito pools. This novel totivirus was recently described in *Cx. inatomii* mosquitoes in Japan (59), and our finding now confirms its association with mosquitoes as a host.

### (+) ssRNA viruses

#### Mesoniviridae

When constructing a phylogenetic tree of the MAG annotated as Alphamesonivirus 1 (99.7 BLASTx % identity; MH520101.1), together with all reference sequences of the *Mesoniviridae* family, our complete Alphamesonivirus Leu4 genome formed a clade with Nam Dinh virus and Cavally virus. Both Alphamesoni 1 viruses are frequently linked to mosquitoes (60, 61). Interestingly, all known members of the *Mesoniviridae* family infect mosquito hosts.

#### Iflaviridae

Iflaviruses are a well-known group of picorna-like viruses that exclusively infect arthropods (62). We found two complete genomes of iflaviruses (Iflavirus Leu5 and Iflavirus Leu6, having a 98.3 and 97.1 BLASTx % identity with Culex iflavi-like virus 4 (MT096522.1) and Yongsan iflavirus 1 (NC_040587.1) respectively) in an urban mosquito pool consisting entirely of *Cx. pipiens* mosquitoes.

#### Negev-related

Negevirus is a proposed taxon for diverse and geographically widely distributed insect-specific viruses isolated from mosquitoes and phlebotomine sandflies (63). We recovered 2 full genomes annotated as Yongsan negev-like virus 1 (average of 95.1 BLASTx % identity; MH703054.1) from two peri-urban mosquito pools that mainly contained *Cx. modestus* mosquitoes, named Negevirus Leu7 and Leu8.

### (-) ssRNA viruses

#### Rhabdoviridae

Rhabdoviruses are a diverse group of negative-sense ssRNA viruses known to infect both vertebrates and invertebrates as well as plants (64). Riversidevirus 1 was first described in *Ochlerotatus sp*. mosquitoes in Central Europe (65) and, in this study, it was also detected (98.2% BLASTx identity; KU248086.1). Rhabdovirus Leu9 was identified in a pool containing mostly *Ochlerotatus* mosquitoes. This suggests a restricted host species range as, up to this date and to our knowledge, this virus has not been found in other mosquito species or other hosts yet.

## Discussion

A national mosquito inventory between 2007 and 2010 (MODIRISK project) clarified that the mosquito fauna in Belgium is composed by twenty-three mosquito species belonging to five traditionally recognized genera, including twenty-one indigenous and two exotic species (*Ae. koreicus* and *Ae. japonicus*)(66). The five most abundant species were *Cx. pipiens* (61.62%), *Coquillettidia richiardii* (15.43%), *Ae. cinereus* (5.04%), *Anopheles claviger* (3.52%) and *Ae. vexans* (2.93%) (66). Amid the eight species that were collected in this study in Leuven, *Cx. pipiens*, *Cx. torrentium*, *Culiseta annulata*, *Culiseta morsitans*, *Ae. sticticus*, *Ae. cinereus* and *Anopheles plumbeus* have been reported as autochthonous species of Belgium according to the latest mosquito species checklist(6).

In contrast, *Cx. modestus* findings in Belgium are rare. Only one larva has been encountered before during the latest exotic mosquito survey carried out from 2017 – 2019 (7, 67) However, during our survey in 2019, *Cx. modestus* accounted for almost half of the mosquitoes that were collected, in three different breeding sites. In addition, *Cx. modestus* mosquitoes were reconfirmed at the same collection sites in the summer of 2020. This finding suggests the establishment of this mosquito species in Belgium, potentially introduced from the UK or Germany. The appearance and spread of *Cx. modestus* in the UK has only been reported recently as well, although this species seems to be abundantly present in certain regions based on recent surveys (2017, 2019) (3). The hypothesis for not noticing its presence in UK before probably relies on the misidentification of *Cx. modestus* by other mosquito species, such as *Cx. torrentium* (3).

Along with the introduction of a new mosquito species in a region, its potential role in the transmission of arboviruses that could cause disease in animals and humans must be evaluated. The presence of *Cx. modestus* in Belgium could be problematic as it is one of most important vectors for *Dirofilaria spp*. such as *Dirofilaria immitis* (68). Furthermore, coexistence of *Cx. pipiens* and *Cx*. Modestus, two important vectors, may increase the risk of transmission for WNV and USUV, given the right circumstances. These two viruses are likely to co-circulate in the same habitat, where birds and *Cx. modestus* mosquitoes play their roles as hosts and vectors, respectively (21). In September 2020, enzootic transmission of WNV in the Netherlands, a neighbouring country of Belgium, was confirmed for the first time by detecting simultaneously the presence of the virus in a local common whitethroat, in field collected mosquito pools and in humans (69). Given the establishment of *Cx. modestus* in Belgium, it would be advisable to implement vector surveillance for this species. In Europe, the higher biting activity displayed by *Cx. modestus* lasts from July until the beginning of the October. However, given the detection of Tahyna virus (an arbovirus) in hibernating *Cx. modestus* mosquitoes in France (14), winter collection can also be considered for the surveillance of mosquito-transmitted pathogens.

In order to examine the genetic structure of the *Cx. modestus* population found in Leuven, we gathered mitochondrial sequences of *Cx. modestus* mosquitoes collected in other countries across Europe and constructed a haplotype network using the MJ method based on 228 partial COI sequences. As recently reported (3), *Cx. modestus* populations across Europe are separated in two lineages. According to this network most Belgian haplotypes were connected to haplotypes from the UK and Germany, suggesting that the mosquito population in Leuven, Belgium could be derived from these two populations. There were three central haplotypes in the lineage II that were shared by several countries. In lineage I, there is one central haplotype that was shared by individuals from Denmark, Spain, and Belgium. This data might indicate that *Cx. modestus* mosquitoes belonging to both lineages are present in Belgium, suggesting the occurrence of at least two independent introduction events.

Vector competence of the mosquito can be influenced by several factors. Bacterial symbionts such as *Wolbachia* have the ability to hinder infection of a variety of pathogens such as chikungunya virus, dengue virus, Zika virus, WNV and malaria-causing *Plasmodium* in different mosquito species (70). It is possible that viral symbionts discovered in mosquitoes may have a similar effect. For instance, the insect-specific virus Nhumirim virus was shown to inhibit the replication of WNV, St Louis encephalitis virus and Japanese encephalitis virus in C6/36 cells (71). As a first step into unveil the role of viral symbionts in the mosquito’s vector competence, we investigated the virome of the collected mosquitoes. Of note, no USUV or WNV was detected in the collected *Cx*. mosquitoes. Furthermore, no Lednice virus was detected in the *Cx. modestus* samples, although this mosquito species was reported to be an important Lednice *Orthobunyavirus* vector (13). In total, 33 eukaryotic viral species could be detected across all our samples in this study, and we recovered 9 (near-)complete genomes of 6 viral species.

When comparing viral hits across the mosquito species and habitat types where they were collected, some similarities could be observed. Mosquito pools belonging to the same genus seemed to have more viruses in common, as shown by the clustering of the *Cx*. mosquito pools or the distinct virome profile presented by the pool composed of *Anopheles/Culiseta* (pool 3) compared to the other pools. Additionally, we observed a clustering of pools per habitat type. In this case, peri-urban mosquito pools harbored several viruses in common, and in great abundance, such as Culex totivirus Leu1, Leu2 and Leu3, and Negevirus Leu7 and Leu8, closely related to Culex inatomii totivirus and Yongsan negev-like virus 1, respectively. Also, the 6 viral species of which the (near-) complete genome was recovered were previously reported as, or clustered together with, viruses associated to mosquitoes, which might hint at the preservation of a core mosquito virome. However, a larger sampling size is needed to suggest that the virome composition and its abundance differ according to genus, local acquisition and ecosystem, and habitat composition.

When comparing our results with a virome study on *Cx. quinquefasciatus* and *Ae. aegypti* mosquitoes collected from Guadeloupe, which is the largest island of the French West Indies in the Caribbean, there were two virus species (Hubei toti-like virus 10 and Hubei partiti-like virus 22) found to be shared with Belgian mosquitoes (72). The fact that the same virus species was found in mosquitoes collected in Belgium and in Guadeloupe could indicate a widespread global movement and/or long host–virus coevolution. Moreover, several viruses were shared with Northern European Swedish mosquitoes (Whidbey virus, Hubei partiti-like virus 22, Chaq virus-like 1, Ista virus, Wuhan Mosquito Virus 6, and Sonnbo virus) (31, 73). At the virus family/order level, the relative virome abundance of the Swedish *Cx. pipiens* was dominated by the Luteo-, Orthomyxo- and Nam Dinh virus. In contrast, the virome of Belgian *Cx. pipiens* was dominated by *Iflaviridae* (pool 2).

When mosquito samples are pooled, as we did in our study, the virome profile could be strongly skewed by one or a few high titer virus infection(s) from a single mosquito in the pool. In a study of Swedish mosquitoes, Pettersson et al. (2019) reported that 30% of all reads of one of the libraries composed of *Cx. torrentium* mosquitoes were annotated to Nam Dinh virus. From pool 1, we recovered the (near-) complete genome of Alphamesonivirus Leu4, which is a member of the *Mesoniviridae* family that contains the Nam Dinh virus. Considering what was reported in Swedish mosquitoes and that pool 1 was the only pool containing one individual of *Cx. torrentium*, we suggest that Alphamesonivirus Leu4 might have been harbored by this mosquito species, as it was not found in any other mosquito pool. In our study, the occurrence of more than one mosquito genera in the same pool was unintentional and resulted from the pooling based on morphological identification. For further research, the use of the individual mosquito body is recommended to perform virome characterization. The feasibility of this approach on single mosquitoes has been evaluated and no significant differences in total reads number and viral reads proportion were found when compared to pooled mosquitoes samples (72).

In conclusion, we here report the establishment of *Cx. modestus* in the surroundings of the city of Leuven, Belgium. The virome of the collected mosquitoes was revealed by a metagenomics approach. As *Cx. modestus* is known to be a vector of WNV, USUV and other arboviruses, surveillance for this mosquito species is recommended.

## Acknowledgements

This project was funded by KU Leuven (C22/18/007 and starting grant STG/19/008). We would like to thank Prof. dr. ir. Raf Aerts and his team at the Division Ecology, Evolution and Biodiversity Conservation, University of Leuven (KU Leuven), for their advice on the selection of mosquito collection sites representing different habitat types. Additionally, we would like to thank Prof. Johan Neyts (KU Leuven) for allowing us the use of his laboratory space and equipment for our experimental work. Equally, we would like to thank Katrien De Wolf from the Unit Entomology, Institute of Tropical Medicine in Belgium for reviewing the manuscript and helpful suggestions.

## References

1. Ficalbi E. 1889. Zanzara di colorito modesto. Boll della Soc Entomol Ital 1:93–94.

2. European Centre for Disease Prevention and Control and European Food Safety Authority. 2020. Culex modestus - current known distribution: May 2020.

3. Hernández-Triana LM, Brugman VA, Pramual P, Barrero E, Nikolova NI, Ruiz-Arrondo I, Kaiser A, Krüger A, Lumley S, Osório HC, Ignjatović-Ćupina A, Petrić D, Laure Setier-Rio M, Bødker R, Johnson N. 2020. Genetic diversity and population structure of Culex modestus across Europe: does recent appearance in the United Kingdom reveal a tendency for geographical spread? Med Vet Entomol 34:86–96.

4. Bødker R, Klitgård K, Byriel DB, Kristensen B. 2014. Establishment of the West Nile virus vector, Culex modestus, in a residential area in Denmark. J Vector Ecol 39:1–3.

5. Lindström A, Lilja T. 2018. First finding of the West Nile virus vector Culex modestus Ficalbi 1889 (Diptera; Culicidae) in Sweden. J Eur Mosq Control Assoc 36.

6. Boukraa S, Dekoninck W, Versteirt V, Schaffner F, Coosemans M, Haubruge E, Francis F. 2015. Updated checklist of the mosquitoes (Diptera: Culicidae) of Belgium. J Vector Ecol 40:398–407.

7. De wolf K, Vanderheyden A, Deblauwe I, Smitz N, Gombeer S, Vanslembrouck a, Meganck K, Dekoninck W, De meyer M, Backeljau T, Müller R, Van bortel W. 2021. First record of the West Nile virus bridge vector *Culex modestus* Ficalbi (Diptera: Culicidae) in Belgium, validated by DNA barcoding. Zootaxa 4920:131–139.

8. Brugman VA, Hernández-Triana LM, England ME, Medlock JM, Mertens PPC, Logan JG, Wilson AJ, Fooks AR, Johnson N, Carpenter S. 2017. Blood-feeding patterns of native mosquitoes and insights into their potential role as pathogen vectors in the Thames estuary region of the United Kingdom. Parasites and Vectors 10:1–12.

9. Brugman VA, England ME, Stoner J, Tugwell L, Harrup LE, Wilson AJ, Medlock JM, Logan JG, Fooks AR, Mertens PPC, Johnson N, Carpenter S. 2017. How often do mosquitoes bite humans in southern England? A standardised summer trial at four sites reveals spatial, temporal and site-related variation in biting rates. Parasites and Vectors 10.

10. Tran A, L’Ambert G, Balança G, Pradier S, Grosbois V, Balenghien T, Baldet T, Lecollinet S, Leblond A, Gaidet-Drapier N. 2017. An Integrative Eco-Epidemiological Analysis of West Nile Virus Transmission. Ecohealth 14:474–489.

11. Ionica AM, Zittra C, Wimmer V, Leitner N, Votýpka J, Modrý D, Mihalca AD, Fuehrer HP. 2017. Mosquitoes in the Danube Delta: Searching for vectors of filarioid helminths and avian malaria. Parasites and Vectors 10.

12. Svobodová M, Volf P, Votýpka J. 2015. Trypanosomatids in ornithophilic bloodsucking Diptera. Med Vet Entomol 29:444–447.

13. Berčič RL, Bányai K, Růžek D, Fehér E, Domán M, Danielová V, Bakonyi T, Nowotny N. 2019. Phylogenetic analysis of lednice Orthobunyavirus. Microorganisms 7.

14. Chippaux A, Rageau J, Mouchet J. 1970. [Hibernation of arbovirus Tahyna in Culex modestus Fic. in France]. C R Acad Sci Hebd Seances Acad Sci D1970/03/23. 270:1648–1650.

15. Vaux AGC, Gibson G, Hernandez-Triana LM, Cheke RA, McCracken F, Jeffries CL, Horton DL, Springate S, Johnson N, Fooks AR, Leach S, Medlock JM. 2015. Enhanced West Nile virus surveillance in the North Kent marshes, UK. Parasites and Vectors 8:91.

16. Monaco F, Goffredo M, Briguglio P, Pinoni C, Polci A, Iannetti S, Pinto S, Marruchella G, Di Francesco G, Di Gennaro A, Pais M, Teodori L, Bruno R, Catalani M, Ruiu A, Lelli R, Savini G. 2015. Descrizione dei focolai di west nile disease nel 2011 nella regione Sardegna, Italia. Vet Ital 51:5–16.

17. Rudolf I, Blažejová H, Šebesta O, Mendel J, Peško J, Betášová L, Straková P, Šikutová S, Hubálek Z. 2018. West Nile virus (lineage 2) in mosquitoes in southern Moravia - awaiting the first autochthonous human cases. Epidemiol Mikrobiol Imunol 67:44–6.

18. Rudolf I, Rettich F, Betášová L, Imrichová K, Mendel J, Hubálek Z, Šikutová S. 2019. West Nile virus (lineage 2) detected for the first time in mosquitoes in Southern Bohemia: new WNV endemic area? Epidemiol Mikrobiol Imunol 68:150–153.

19. Balenghien T, Vazeille M, Reiter P, Schaffner F, Zeller H, Bicout D. 2007. Evidence of Laboratory Vector Competence of Culex Modestus for West Nile Virus. J Am Mosq Control Assoc 23.

20. Balenghien T, Vazeille M, Grandadam M, Schaffner F, Zeller H, Reiter P, Sabatier P, Fouque F, Bicout DJ. 2008. Vector competence of some French Culex and Aedes mosquitoes for West Nile Virus. Vector-Borne Zoonotic Dis 8:589–595.

21. Rudolf I, Bakonyi T, Šebesta O, Mendel J, Peško J, Betášová L, Blažejová H, Venclíková K, Straková P, Nowotny N, Hubálek Z. 2015. Co-circulation of Usutu virus and West Nile virus in a reed bed ecosystem. Parasites and Vectors 8.

22. Gaibani P, Rossini G. 2017. An overview of Usutu virus. Microbes Infect 19:382–387.

23. Weissenböck H, Bakonyi T, Rossi G, Mani P, Nowotny N. 2013. Usutu virus, Italy, 1996. Emerg Infect Dis 19:274–277.

24. Weissenböck H, Kolodziejek J, Url A, Lussy H, Rebel-Bauder B, Nowotny N. 2002. Emergence of *Usutu virus*, an African Mosquito-Borne *Flavivirus* of the Japanese Encephalitis Virus Group, Central Europe. Emerg Infect Dis 8:652–656.

25. Becker N, Jöst H, Ziegler U, Eiden M, Höper D, Emmerich P, Fichet-Calvet E, Ehichioya DU, Czajka C, Gabriel M, Hoffmann B, Beer M, Tenner-Racz K, Racz P, Günther S, Wink M, Bosch S, Konrad A, Pfeffer M, Groschup MH, Schmidt-Chanasit J. 2012. Epizootic emergence of Usutu virus in wild and captive birds in Germany. PLoS One 7.

26. Oude Munnink BB, Münger E, Nieuwenhuijse DF, Kohl R, van der Linden A, Schapendonk CME, van der Jeugd H, Kik M, Rijks JM, Reusken CBEM, Koopmans M. 2020. Genomic monitoring to understand the emergence and spread of Usutu virus in the Netherlands, 2016-2018. Sci Rep 10:2798.

27. Rouffaer LO, Steensels M, Verlinden M, Vervaeke M, Boonyarittichaikij R, Martel A, Lambrecht B. 2018. Usutu Virus Epizootic and *Plasmodium* Coinfection in Eurasian Blackbirds (*Turdus merula*) in Flanders, Belgium. J Wildl Dis 54:859–862.

28. Benzarti E, Sarlet M, Franssen M, Cadar D, Schmidt-Chanasit J, Rivas JF, Linden A, Desmecht D, Garigliany M. 2020. Usutu Virus Epizootic in Belgium in 2017 and 2018: Evidence of Virus Endemization and Ongoing Introduction Events. Vector-Borne Zoonotic Dis 20:43–50.

29. Shi C, Liu Y, Hu X, Xiong J, Zhang B, Yuan Z. 2015. A metagenomic survey of viral abundance and diversity in mosquitoes from hubei province. PLoS One. Public Library of Science.

30. Bolling BG, Weaver SC, Tesh RB, Vasilakis N. 2015. Insect-specific virus discovery: Significance for the arbovirus community. Viruses. MDPI AG.

31. Öhlund P, Hayer J, Lundén H, Hesson JC, Blomström A-L. 2019. Viromics Reveal a Number of Novel RNA Viruses in Swedish Mosquitoes. Viruses 11:1027.

32. Mayi MPA, Bamou R, Djiappi-Tchamen B, Fontaine A, Jeffries CL, Walker T, Antonio-Nkondjio C, Cornel AJ, Tchuinkam T. 2020. Habitat and Seasonality Affect Mosquito Community Composition in the West Region of Cameroon. Insects 11:312.

33. Becker N, Petric D, Zgomba M, Boase C, Madon M, Dahl C, Kaiser A, Becker N, Petrić D, Zgomba M, Boase C, Madon M, Dahl C, Kaiser A. 2010. Key to Female Mosquitoes, p. 91–111. In Mosquitoes and Their Control. Springer Berlin Heidelberg.

34. Ander M, Troell K, Chirico J. 2013. Barcoding of biting midges in the genus Culicoides: A tool for species determination. Med Vet Entomol 27:323–331.

35. Folmer O, Black M, Hoeh W, Lutz R, Vrijenhoek R. 1994. DNA primers for amplification of mitochondrial cytochrome c oxidase subunit I from diverse metazoan invertebrates. Mol Mar Biol Biotechnol 3:294–299.

36. Hall TA. 1999. BioEdit: a user-friendly biological sequence alignment editor and analysis program for Windows 95/98/NT. Nucleic Acids Symp Ser 41:95–98.

37. Versteirt V, Nagy ZT, Roelants P, Denis L, Breman FC, Damiens D, Dekoninck W, Backeljau T, Coosemans M, Van Bortel W. 2015. Identification of Belgian mosquito species (Diptera: Culicidae) by DNA barcoding. Mol Ecol Resour 15:449–457.

38. Katoh K, Standley DM. 2013. MAFFT multiple sequence alignment software version 7: Improvements in performance and usability. Mol Biol Evol 30:772–780.

39. Capella-Gutiérrez S, Silla-Martínez JM, Gabaldón T. 2009. trimAl: A tool for automated alignment trimming in large-scale phylogenetic analyses. Bioinformatics 25:1972–1973.

40. Criscuolo A, Gribaldo S. 2010. BMGE (Block Mapping and Gathering with Entropy): A new software for selection of phylogenetic informative regions from multiple sequence alignments. BMC Evol Biol 10:210.

41. Nguyen LT, Schmidt HA, Von Haeseler A, Minh BQ. 2015. IQ-TREE: A fast and effective stochastic algorithm for estimating maximum-likelihood phylogenies. Mol Biol Evol 32:268–274.

42. Excoffier L, Lischer HEL. 2010. Arlequin suite ver 3.5: A new series of programs to perform population genetics analyses under Linux and Windows. Mol Ecol Resour 10:564–567.

43. Bandelt HJ, Forster P, Röhl A. 1999. Median-joining networks for inferring intraspecific phylogenies. Mol Biol Evol 16:37–48.

44. Leigh JW, Bryant D. 2015. POPART: Full-feature software for haplotype network construction. Methods Ecol Evol 6:1110–1116.

45. Conceição-Neto N, Zeller M, Lefrère H, De Bruyn P, Beller L, Deboutte W, Yinda CK, Lavigne R, Maes P, Ranst M Van, Heylen E, Matthijnssens J. 2015. Modular approach to customise sample preparation procedures for viral metagenomics: A reproducible protocol for virome analysis. Sci Rep 5:1–14.

46. Shi C. 2020. Unbiased analyses of virome in mosquito vectors and their association with the transmission potential of pathogenetic arboviruses. KU Leuven.

47. Bolger AM, Lohse M, Usadel B. 2014. Trimmomatic: A flexible trimmer for Illumina sequence data. Bioinformatics 30:2114–2120.

48. Langmead B, Salzberg SL. 2012. Fast gapped-read alignment with Bowtie 2. Nat Methods 9:357–359.

49. Nurk S, Meleshko D, Korobeynikov A, Pevzner PA. 2017. MetaSPAdes: A new versatile metagenomic assembler. Genome Res 27:824–834.

50. Buchfink B, Xie C, Huson DH. 2015. Fast and sensitive protein alignment using DIAMOND. Nat Methods. Nature Publishing Group.

51. Ondov BD, Bergman NH, Phillippy AM. 2011. Interactive metagenomic visualization in a Web browser. BMC Bioinformatics 12:385.

52. Li H, Durbin R. 2009. Fast and accurate short read alignment with Burrows-Wheeler transform. Bioinformatics 25:1754–1760.

53. Md V, Misra S, Li H, Aluru S. 2019. Efficient Architecture-Aware Acceleration of BWA-MEM for Multicore Systems. IEEE Parallel Distrib Process Symp 314–324.

54. McMurdie PJ, Holmes S. 2013. phyloseq: An R Package for Reproducible Interactive Analysis and Graphics of Microbiome Census Data. PLoS One 8:e61217.

55. Paulson JN, Colin Stine O, Bravo HC, Pop M. 2013. Differential abundance analysis for microbial marker-gene surveys. Nat Methods 10:1200–1202.

56. Dixon P. 2003. VEGAN, a package of R functions for community ecology. J Veg Sci. Opulus Press AB.

57. Gu Z, Eils R, Schlesner M. 2016. Complex heatmaps reveal patterns and correlations in multidimensional genomic data. Bioinformatics 32:2847–2849.

58. Li H, Handsaker B, Wysoker A, Fennell T, Ruan J, Homer N, Marth G, Abecasis G, Durbin R. 2009. The Sequence Alignment/Map format and SAMtools. Bioinformatics 25:2078–2079.

59. Faizah AN, Kobayashi D, Isawa H, Amoa-Bosompem M, Murota K, Higa Y, Futami K, Shimada S, Kim KS, Itokawa K, Watanabe M, Tsuda Y, Minakawa N, Miura K, Hirayama K, Sawabe K. 2020. Deciphering the Virome of Culex vishnui Subgroup Mosquitoes, the Major Vectors of Japanese Encephalitis, in Japan. Viruses 12:264.

60. Zhou J, Jin Y, Chen Y, Li J, Zhang Q, Xie X, Gan L, Liu Q. 2017. Complete Genomic and Ultrastructural Analysis of a Nam Dinh Virus Isolated from Culex pipiens quinquefasciatus in China^*^. Sci Rep 7:271.

61. Zirkel F, Kurth A, Quan PL, Briese T, Ellerbrok H, Pauli G, Leendertz FH, Lipkin WI, Ziebuhr J, Drosten C, Junglen S. 2011. An insect nidovirus emerging from a primary tropical rainforest. MBio 2.

62. van Oers M. 2008. Iflavirus, p. 42–6. In Mahy, B, Van Regenmortel, M (eds.), Encyclopedia of VirologyThird. Oxford: Academic Press.

63. Vasilakis N, Forrester NL, Palacios G, Nasar F, Savji N, Rossi SL, Guzman H, Wood TG, Popov V, Gorchakov R, Gonzalez A V., Haddow AD, Watts DM, da Rosa APAT, Weaver SC, Lipkin WI, Tesh RB. 2013. Negevirus: a Proposed New Taxon of Insect-Specific Viruses with Wide Geographic Distribution. J Virol 87:2475–2488.

64. Kuzmin I V., Novella IS, Dietzgen RG, Padhi A, Rupprecht CE. 2009. The rhabdoviruses: Biodiversity, phylogenetics, and evolution. Infect Genet Evol 9:541–553.

65. Reuter G, Boros Á, Pál J, Kapusinszky B, Delwart E, Pankovics P. 2016. Detection and genome analysis of a novel (dima)rhabdovirus (Riverside virus) from Ochlerotatus sp. mosquitoes in Central Europe. Infect Genet Evol 39:336–341.

66. Versteirt V, Boyer S, Damiens D, De Clercq EM, Dekoninck W, Ducheyne E, Grootaert P, Garros C, Hance T, Hendrickx G, Coosemans M, Van Bortel W. 2013. Nationwide inventory of mosquito biodiversity (Diptera: Culicidae) in Belgium, Europe. Bull Entomol Res 103:193–203.

67. Deblauwe I, De Wolf K, Smitz N, Vanslembrouck A, Schneider A, De Witte J, Verlé I, Dekoninck W, De Meyer M, Backeljau T, Gombeer S, Meganck K, Van Bourgonie Y-R, Vanderheyden A, Müller R, Van Bortel W. 2020. Monitoring of exotic mosquitoes in Belgium (MEMO): Final Report Phase 7 Part 1: MEMO results. Antwerp.

68. Rossi L, Pollono F, Meneguz PG, Cancrini G. 1999. Quattro specie di culicidi come possibili vettori di Dirofilaria immitis nella risaia piemontese. Parassitologia 41:537–542.

69. Vlaskamp DR, Thijsen SF, Reimerink J, Hilkens P, Bouvy WH, Bantjes SE, Vlaminckx BJ, Zaaijer H, van den Kerkhof HH, Raven SF, Reusken CB. 2020. First autochthonous human West Nile virus infections in the Netherlands, July to August 2020. Eurosurveillance 25:2001904.

70. Caragata EP, Dutra HLC, Moreira LA. 2016. Inhibition of Zika virus by Wolbachia in Aedes aegypti. Microb Cell. Shared Science Publishers OG.

71. Kenney JL, Solberg OD, Langevin SA, Brault AC. 2014. Characterization of a novel insect-specific flavivirus from Brazil: Potential for inhibition of infection of arthropod cells with medically important flaviviruses. J Gen Virol 95:2796–2808.

72. Shi C, Beller L, Deboutte W, Yinda KC, Delang L, Vega-Rúa A, Failloux AB, Matthijnssens J. 2019. Stable distinct core eukaryotic viromes in different mosquito species from Guadeloupe, using single mosquito viral metagenomics. Microbiome 7:121.

73. Pettersson, Shi, Eden, Holmes, Hesson. 2019. Meta-Transcriptomic Comparison of the RNA Viromes of the Mosquito Vectors Culex pipiens and Culex torrentium in Northern Europe. Viruses 11:1033.

