## Supplementary materials for "Establishment of *Culex modestus* in Belgium and a glance into the virome of Belgian mosquito species"

### **Supplementary material**

#### **Figure S1. Overview of the collection sites and points in Leuven.** Overview of collection sites representing three distinct habitat types in Leuven: Botanic Garden (Urban), Arenberg Castle (Peri-urban), and Wetlands (Source: Google Maps). A BioGent Sentinel trap in collection site is displayed on the bottom right corner.

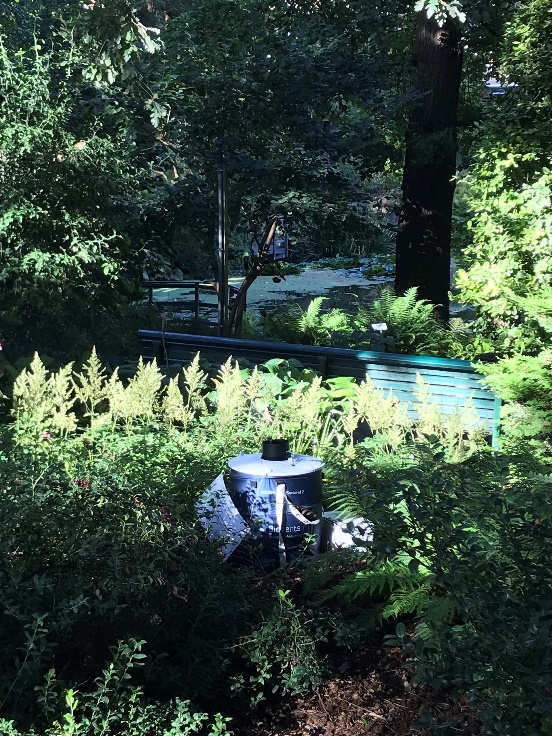

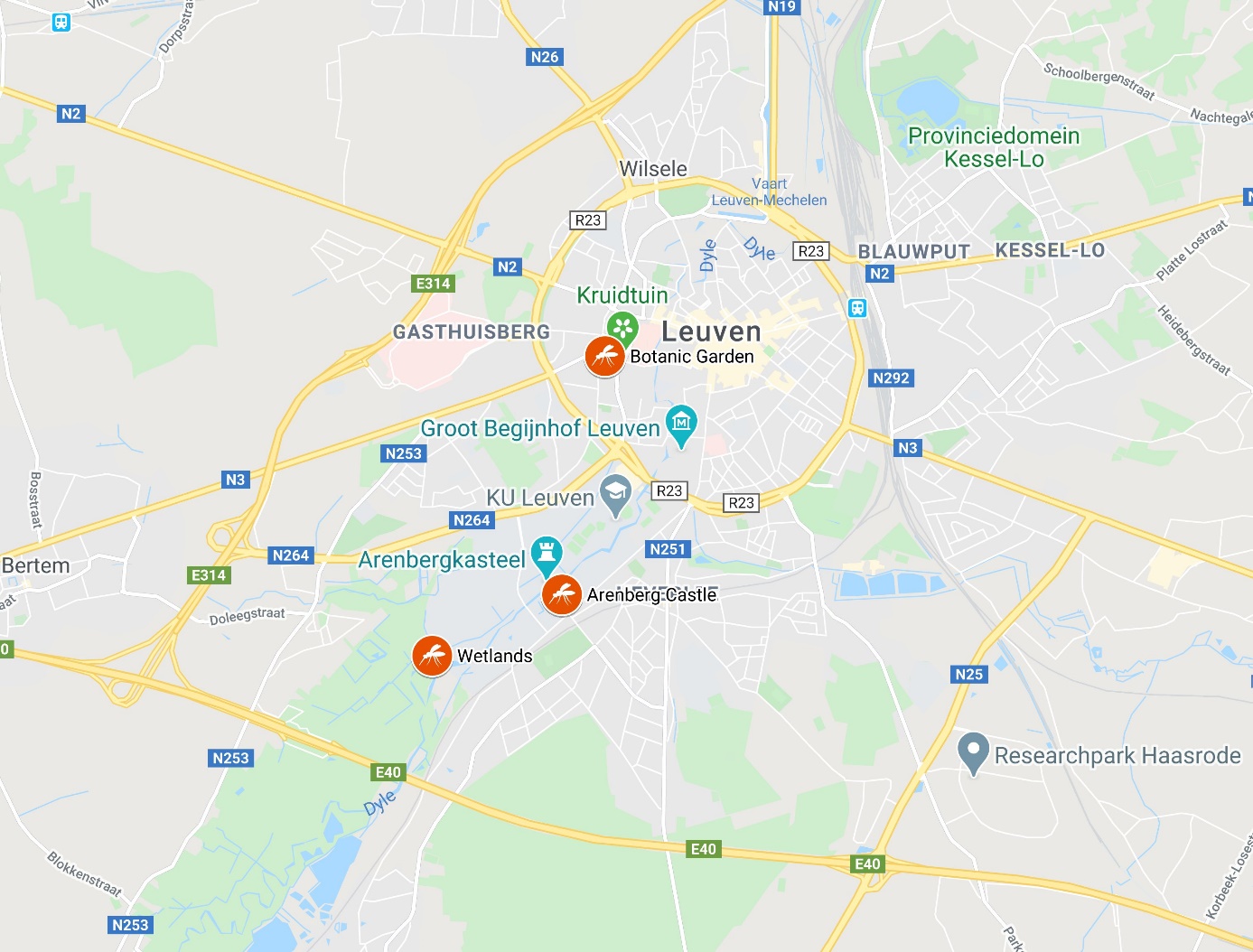

#### **Table S2. *Culex modestus* sequences downloaded from Genbank and employed for the construction of the haplotype network.**

| **Country** | **Locality** | **Accession number (Genbank)** |
| --- | --- | --- |
| Spain | Calahorra | MK402890, MK402903 |
|  | Viana | MK402912, MK971866, MK971936, MK971952, MK971972 |
|  | Hervias | MK402733, MK402793, MK402814, MK402818, MK402875, MK402881, MK402897, MK971950 |
|  | Haro | MK402688, MK402692, MK402724, MK402798 |
|  | Logrono | MK402689, MK402754, MK402845 |
| Germany | Speyer | MK971796, MK971801, MK971805, MK971807, MK971814, MK971863, MK971869, MK971872, MK971880, MK971911, MK971914, MK971923, MK971931, MK971963, MK971975, MK971989, MK971991, MK972008 |
|  | Roememberg | MK971808, MK971828, MK971848, MK971857, MK971861, MK971876, MK971881, MK971902, MK971909, MK971960, MK972005 |
|  | Waghausel | MK971883, MK971980 |
|  | Ketsch | MK971940 |
|  | Trebur | MK971885 |
|  | Lingenfeld | MK971824, MK971826, MK971832, MK971858, MK971939, MK971957, MK971983 |
|  | Frankfurt | HF562836, HF562837 |
| United Kingdom | Cliffe | JN592733, JN592734, JN592735, MK971798, MK971799, MK971803, MK971809, MK971818, MK971835, MK971845, MK971847, MK971853, MK971860, MK971865, MK971875, MK971878, MK971887, MK971889, MK971893, MK971896, MK971899, MK971905, MK971910, MK971916, MK971920, MK971926, MK971927, MK971935, MK971941, MK971954, MK971966, MK971968, MK971970, MK971982, MK971999, MK972011 |
|  | Elmley | MK971995, JN592729, JN592730, KU877022, MK403130, MK403139, MK403162, MK403205, MK403242, MK403257, MK403293, MK403339, MK403462, MK403478, MK971797, MK971819, MK971821, MK971825, MK971827, MK971829, MK971844, MK971846, MK971854, MK971891, MK971904, MK971915, MK971934, MK971942, MK971949, MK971976, MK971994, MK972001, MK972003 |
|  | Northward Hill | JN592742, JN592743, JN592744, JN592745 |
|  | Kent | MK403327 |
| France | Arles | MK971812, MK971823, MK971833, MK971834, MK971839, MK971852, MK971856, MK971867, MK971870, MK971871, MK971882, MK971903, MK971937, MK971943, MK971947, MK971951, MK971958, MK971967, MK971971, MK971978, MK971981, MK971984, MK971990, MK971996, MK972002 |
|  | Camargue | JN592746, JN592747, JN592748 |
| Serbia | Novid sad | MK971874, MK971884, MK971961, MK971998 |
| Portugal | Albufeira | MK971810 |
|  | Alcacer | MK971924 |
| Denmark | Greve | KJ401301, KJ401302, KJ401303, KJ401304, KJ401305 |
|  | Copenhagen | MK971877, MK971890, |
| Sweden | Falsterbo | MG844175, MG844176, MG844178 |
|  | Simrishamn | MF537266, MG844177 |

#### **Table S3. Haplotype frequencies among populations of *Culex modestus* from 9 countries in Europe.**

|  | **Belgium** | **Denmark** | **France** | **Germany** | **Portugal** | **Serbia** | **Spain** | **Sweden** | **UK** |
| --- | --- | --- | --- | --- | --- | --- | --- | --- | --- |
| Hap_1 | 1 |  |  |  |  |  |  |  |  |
| Hap_2 | 1 |  |  |  |  |  |  |  |  |
| Hap_3 | 1 |  |  |  |  |  |  |  |  |
| Hap_4 | 3 |  |  |  |  |  |  |  |  |
| Hap_5 | 1 |  |  |  |  |  |  |  |  |
| Hap_6 | 1 |  |  |  |  |  |  |  |  |
| Hap_7 | 1 |  |  |  |  |  |  |  |  |
| Hap_8 | 1 | 1 | 3 |  |  |  | 3 |  |  |
| Hap_9 | 2 |  |  |  |  |  |  |  |  |
| Hap_10 | 1 |  |  |  |  |  |  |  |  |
| Hap_11 | 1 |  |  |  |  |  |  |  |  |
| Hap_12 | 3 |  | 2 | 2 |  |  |  | 1 | 2 |
| Hap_13 | 1 |  |  |  |  |  |  |  |  |
| Hap_14 | 2 |  |  |  |  |  |  |  |  |
| Hap_15 | 1 |  |  |  |  |  |  |  |  |
| Hap_16 | 1 |  |  |  |  |  |  |  |  |
| Hap_17 | 1 |  |  |  |  |  |  |  |  |
| Hap_18 | 2 |  |  |  |  |  |  |  |  |
| Hap_19 | 1 |  |  |  |  |  |  |  |  |
| Hap_20 | 3 |  |  | 1 |  |  |  |  | 1 |
| Hap_21 | 1 |  |  |  |  |  |  |  |  |
| Hap_22 | 1 |  |  |  |  |  |  |  |  |
| Hap_23 | 1 |  |  |  |  |  |  |  |  |
| Hap_24 | 2 |  |  |  |  |  |  |  |  |
| Hap_25 | 1 |  |  |  |  |  |  | 1 | 2 |
| Hap_26 | 1 |  |  |  |  |  |  |  |  |
| Hap_27 | 2 |  |  |  |  | 1 |  |  | 2 |
| Hap_28 | 1 |  |  |  |  |  |  |  |  |
| Hap_29 | 1 |  |  |  |  |  |  |  |  |
| Hap_30 | 1 | 1 |  |  |  |  |  | 1 |  |
| Hap_31 | 1 |  |  |  |  |  |  |  |  |
| Hap_32 | 1 |  |  |  |  |  |  |  |  |
| Hap_33 | 1 |  |  |  |  |  |  |  |  |
| Hap_34 |  | 1 |  |  |  |  |  | 1 |  |
| Hap_35 |  | 1 |  |  |  |  |  |  |  |
| Hap_36 |  | 1 |  | 2 |  |  |  |  |  |
| Hap_37 |  | 1 |  |  |  |  |  |  |  |
| Hap_38 |  | 1 |  |  |  |  |  |  |  |
| Hap_39 |  |  | 1 |  |  |  |  |  |  |
| Hap_40 |  |  | 9 |  |  |  |  |  |  |
| Hap_41 |  |  | 1 |  |  |  |  |  |  |
| Hap_42 |  |  | 1 |  |  |  |  |  |  |
| Hap_43 |  |  | 5 |  |  |  |  |  |  |
| Hap_44 |  |  | 3 |  |  |  |  |  |  |
| Hap_45 |  |  | 1 |  |  |  |  |  | 3 |
| Hap_46 |  |  | 1 |  |  |  |  |  |  |
| Hap_47 |  |  | 1 |  |  |  |  |  |  |
| Hap_48 |  |  |  | 9 |  |  |  |  |  |
| Hap_49 |  |  |  | 9 |  |  |  |  |  |
| Hap_50 |  |  |  | 2 |  |  |  |  |  |
| Hap_51 |  |  |  | 3 |  |  |  |  |  |
| Hap_52 |  |  |  | 1 |  |  |  |  |  |
| Hap_53 |  |  |  | 4 |  |  |  |  |  |
| Hap_54 |  |  |  | 1 |  |  |  |  |  |
| Hap_55 |  |  |  | 1 |  |  |  |  |  |
| Hap_56 |  |  |  | 1 |  |  |  |  |  |
| Hap_57 |  |  |  | 1 |  |  |  |  |  |
| Hap_58 |  |  |  | 2 |  |  |  |  |  |
| Hap_59 |  |  |  | 1 |  |  |  |  |  |
| Hap_60 |  |  |  | 1 |  |  |  |  |  |
| Hap_61 |  |  |  | 1 |  |  |  |  |  |
| Hap_62 |  |  |  |  | 1 |  |  |  |  |
| Hap_63 |  |  |  |  | 1 |  |  |  |  |
| Hap_64 |  |  |  |  |  | 1 |  |  |  |
| Hap_65 |  |  |  |  |  | 1 |  |  |  |
| Hap_66 |  |  |  |  |  | 1 |  |  |  |
| Hap_67 |  |  |  |  |  |  | 2 |  |  |
| Hap_68 |  |  |  |  |  |  | 5 |  |  |
| Hap_69 |  |  |  |  |  |  | 1 |  |  |
| Hap_70 |  |  |  |  |  |  | 8 |  |  |
| Hap_71 |  |  |  |  |  |  | 1 |  |  |
| Hap_72 |  |  |  |  |  |  | 1 |  |  |
| Hap_73 |  |  |  |  |  |  | 1 |  |  |
| Hap_74 |  |  |  |  |  |  |  | 1 |  |
| Hap_75 |  |  |  |  |  |  |  |  | 1 |
| Hap_76 |  |  |  |  |  |  |  |  | 15 |
| Hap_77 |  |  |  |  |  |  |  |  | 4 |
| Hap_78 |  |  |  |  |  |  |  |  | 1 |
| Hap_79 |  |  |  |  |  |  |  |  | 5 |
| Hap_80 |  |  |  |  |  |  |  |  | 1 |
| Hap_81 |  |  |  |  |  |  |  |  | 1 |
| Hap_82 |  |  |  |  |  |  |  |  | 2 |
| Hap_83 |  |  |  |  |  |  |  |  | 11 |
| Hap_84 |  |  |  |  |  |  |  |  | 6 |
| Hap_85 |  |  |  |  |  |  |  |  | 1 |
| Hap_86 |  |  |  |  |  |  |  |  | 1 |
| Hap_87 |  |  |  |  |  |  |  |  | 1 |
| Hap_88 |  |  |  |  |  |  |  |  | 1 |
| Hap_89 |  |  |  |  |  |  |  |  | 1 |
| Hap_90 |  |  |  |  |  |  |  |  | 2 |
| Hap_91 |  |  |  |  |  |  |  |  | 2 |
| Hap_92 |  |  |  |  |  |  |  |  | 1 |
| Hap_93 |  |  |  |  |  |  |  |  | 2 |
| Hap_94 |  |  |  |  |  |  |  |  | 1 |
| Hap_95 |  |  |  |  |  |  |  |  | 2 |
| Hap_96 |  |  |  |  |  |  |  |  | 1 |
| Hap_97 |  |  |  |  |  |  |  |  | 1 |

#### **Table S3. Haplotypes among populations of *Culex modestus* from 9 countries in Europe.**

| **Country** | **Locality** | **Haplotypes (n)** |
| --- | --- | --- |
| Belgium (2019) | Leuven | Hap1, Hap2, Hap3, Hap4(3), Hap5, Hap7, Hap8, Hap9(2), Hap10, Hap11, Hap12(2), Hap13, Hap14(2), Hap15, Hap16, Hap17, Hap18(2), Hap20(2), Hap22, Hap23, Hap24(2), Hap25, Hap26, Hap27(2), Hap28, Hap29, Hap30, Hap31, Hap32, Hap33, |
| Belgium (2020) | Leuven | Hap12, Hap19, Hap20, Hap21 |
| Spain | Calahorra | Hap8, Hap68, |
|  | Viana | Hap68(2), Hap70(4), |
|  | Hervias | Hap8, Hap67, Hap70(2), Hap72, Hap73, |
|  | Haro | Hap8, Hap67, Hap69, Hap70(2), |
|  | Logrono | Hap68(2), Hap71, |
| Germany | Speyer | Hap12(2), Hap36, Hap48(4), Hap49(2), Hap50(2), Hap53, Hap54, Hap56, Hap57, Hap58, Hap60, Hap61, |
|  | Roememberg | Hap20, Hap48, Hap49(3), Hap51(3), Hap53(2), Hap59, |
|  | Waghausel | Hap49, Hap55, |
|  | Ketsch | Hap49, |
|  | Trebur | Hap49, |
|  | Lingenfeld | Hap36, Hap48(2), Hap49, Hap52, Hap53, Hap58, |
|  | Frankfurt | Hap48(2), |
| United Kingdom | Cliffe | Hap12(2), Hap25, Hap27, Hap45, Hap76(5), Hap77(3), Hap78, Hap79(2), Hap83(7), Hap84(4), Hap90(2), Hap91(2), Hap93(2), Hap94, Hap96, Hap97 |
|  | Elmley | Hap20, Hap25, Hap27, Hap45(2), Hap75, Hap76(10), Hap77, Hap79(2), Hap82, Hap83(4), Hap84(2), Hap85, Hap86, Hap87, Hap89, Hap92, Hap95(2) |
|  | Northward Hill | Hap79, Hap80, Hap81, Hap82, |
|  | Kent | Hap88 |
| France | Arles | Hap8(3), Hap12(2), Hap40(8), Hap42, Hap43(5), Hap44(3), Hap45, Hap46, Hap47, |
|  | Camargue | Hap39, Hap40, Hap41, |
| Serbia | Novid sad | Hap27, Hap64, Hap65, Hap66 |
| Portugal | Albufeira | Hap62 |
|  | Alcacer | Hap63 |
| Denmark | Greve | Hap8, Hap30, Hap34, Hap35, Hap36, |
|  | Copenhagen | Hap37, Hap38, |
| Sweden | Falsterbo | Hap12, Hap34, Hap74, |
|  | Simrishamn | Hap25, Hap30, |

#### **Table S4. Summary information of the complete genomes identified in this study.**

| **Virus** | **Closest match (Blastx)** | **Accession number** | **% Per. Identity**  **(aa)** | **Genome size (bp)** |
| --- | --- | --- | --- | --- |
| Culex totivirus Leu1 | Culex inatomii totivirus (capsid protein) | LC514398.1 | 98,3 | 6241 |
| Culex totivirus Leu2 | Culex inatomii totivirus (capsid protein) | LC514398.1 | 98,3 | 6273 |
| Culex totivirus Leu3 | Culex inatomii totivirus (capsid protein) | LC514398.1 | 98,2 | 6200 |
| Alphamesonivirus Leu 4 | Alphamesonivirus 1 (pp1ab polyprotein) | MH520101.1 | 99,7 | 20153 |
| Iflavirus Leu5 | Culex iflavi-like virus 4 (polyprotein) | MT096522.1 | 98,3 | 9634 |
| Iflavirus Leu6 | Yongsan iflavirus 1 (polyprotein) | NC_040587.1 | 97,1 | 8977 |
| Negevirus Leu7 | Yongsan negev-like virus 1 (RdRp) | MH703054.1 | 94,6 | 8388 |
| Negevirus Leu8 | Yongsan negev-like virus 1 (RdRp) | MH703054.1 | 95,6 | 8312 |
| Rhabdovirus Leu9 | Riverside virus 1 (large protein) | KU248086.1 | 98,2 | 11659 |
